# Hypoxia-induced switch in SNAT2/SLC38A2 regulation generates endocrine-resistance in breast cancer

**DOI:** 10.1101/384628

**Authors:** Matteo Morotti, Esther Bridges, Alessandro Valli, Hani Choudhry, Helen Sheldon, Simon Wigfield, Nicki Gray, Dylan Jones, Eugene J. Teoh, Wei-Chen Cheng, Simon Lord, Syed Haider, Alan McIntyre, Deborah C. I. Goberdhan, Francesca Buffa, Adrian L. Harris

**Author notes:** **Corresponding author:** Adrian L. Harris Hypoxia and Angiogenesis Group, Cancer Research UK Molecular Oncology Laboratories, Weatherall Institute of Molecular Medicine, University of Oxford, Oxford OX3 9DS, United Kingdom.; Matteo Morotti Hypoxia and Angiogenesis Group, Cancer Research UK Molecular Oncology Laboratories, Weatherall Institute of Molecular Medicine, University of Oxford, Oxford OX3 9DS, United Kingdom.

## Abstract

Tumor hypoxia is associated with poor patient outcomes in estrogen receptor-α (ERα) positive breast cancer. Hypoxia is known to affect tumor growth by reprogramming metabolism and regulating amino acid (AA) uptake. Here we show that the glutamine transporter, SNAT2, is the AA transporter most frequently induced by hypoxia in breast cancer and it is regulated by HIF1α both *in-vitro* and *in-vivo* in xenografts. SNAT2 induction in MCF7 cells was also regulated by ERα but it became predominantly a HIF-1α-dependent gene under hypoxia. Relevant to this, binding sites for both HIF-1α and ERα overlap in SNAT2’s cis-regulatory elements. In addition, the downregulation of SNAT2 by the ER antagonist fulvestrant was reverted in hypoxia.

Overexpression of SNAT2 *in-vitro* to recapitulate the levels induced by hypoxia caused enhanced growth, particularly after ERα inhibition, in hypoxia, or when glutamine levels were low. SNAT2 upregulation *in-vivo* caused complete resistance to anti-estrogen and, partially, anti-VEGF therapies. Finally, high SNAT2 expression levels correlate with HIF-1α and worse outcome in patients given anti-estrogen therapy. Our findings show a switch in regulation of SNAT2 between ERα and HIF-1α, leading to endocrine resistance in hypoxia. Development of drugs targeting SNAT2 may be of value for a subset of hormone-resistant breast cancer.

## Introduction

The estrogen receptor positive (ERα+) subtype accounts for approximately 70% of all newly diagnosed cases of breast cancer in Europe and the USA (1). The majority of these tumors depend on estrogen signaling, thereby providing the rationale for using anti-estrogens as adjuvant therapy to treat breast cancer. Endocrine agents targeting estrogen receptor alpha (ERα) such as tamoxifen, fulvestrant/faslodex or inhibiting estrogen biosynthesis, such as aromatase inhibitors, represent the cornerstone of systemic treatment of this breast cancer subtype, for both early and metastatic disease (2). However, despite recent therapeutic advances, poor response and resistance limit the effectiveness of these agents in up to 30 *%* of patients, manifesting as either relapse after therapy or as disease progression in the metastatic setting (3).

Recent randomized trials confirmed that first line endocrine therapy with Fulvestrant alone or in combination with palbociclib, a Cyclin-dependent kinase 4 and 6 inhibitor, improve clinical outcomes in ER+ metastatic breast cancer patients with disease progression or early relapse following anti-estrogen treatment (4-6).

However, as seen for other hormonal treatments, resistance eventually develops and leads to disease progression.

Many mechanisms have been proposed to account for endocrine resistance such as genetic (loss of ERα expression, mutation, expression of truncated ER isoforms), post-translational modification of ERα and activation by peptide growth factors and epigenetic (methylation, promoter inhibition) changes within the tumor that activate hormone-independent mitogenic pathways (7). In addition to cancer cell-autonomous factors, the host microenvironment can contribute to endocrine resistance (8).

Hypoxia is a key physiological and microenvironmental difference between tumor and normal tissues and is related to poor clinical prognosis and resistance to therapies in many solid tumors, including breast cancer (9, 10). Indeed hypoxia is associated with resistance to chemoendocrine treatment in breast cancer patients (11).

Furthermore, increased expression of hypoxia-inducible factor 1α (HIF-1α), the key transcription factor mediating hypoxia response, is associated with tamoxifen resistance in neoadjuvant, primary therapy of ERα+ breast cancers (12) and is associated with resistance to primary endocrine therapy in primary ER+ breast cancers (13). Moreover, we recently found that the *HIF-1α* gene bears a canonical ER-binding element that responds to estrogen signaling, demonstrating a direct regulatory link between the ERα and HIF-1α pathways in breast cancer (14). HIF-1α function conferred resistance to fulvestrant and could therefore compensate for estrogen signaling when ERα function is compromised during hormone therapy, and thus cause resistance to tamoxifen or fulvestrant treatment (14).

A crucial mechanism by which the growth of cancer cells is promoted in hypoxic microenvironments is by metabolic reprogramming (15). The adaptation to hypoxia is coordinated by HIF-1α, which induces metabolic genes involved in increasing glycolytic flux (16). There is also upregulation of glutamine metabolism to support proliferation, lipid biosynthesis and protection from free radical stress (17, 18). In addition to pyruvate derived from glycolysis, hypoxic cancer cells can supply substrates to the tricarboxylic acid (TCA) cycle to sustain mitochondrial ATP production (anaplerosis) through the uptake of amino acids (AAs), such as glutamine, glycine and serine (19). In particular glutamine, the most abundant AA in the plasma, can fuel the TCA cycle through a series of biochemical reactions termed glutaminolysis (20).

This gives a strong rationale to identify hypoxia-induced metabolic alterations, particularly regarding glutaminolysis. These have the potential to provide induced essentiality opportunities in combination with other drugs (21). Metabolomic analysis of breast cancer patients showed that cancer cells have increased glutamine metabolism compared to normal cells. Compared with the ratio of glutamate to glutamine (indirect assessment of glutaminolysis) in normal tissues, 56% of the ER+ tumor tissues were glutamate enriched (22). Several studies have shown that endocrine therapy resistance in breast cancer cells is modulated by metabolic rewiring and tamoxifen-resistant cells are characterized by HIF-1α hyper-activation via modulation of Akt/mTOR, thus resulting in enhanced aerobic glycolysis and mitochondrial metabolism (23, 24). These data highlight the importance of metabolic adaptability of cancer cells for endocrine therapy resistance and suggest that targeting glutamine metabolism could be a novel approach to overcome resistance to endocrine therapies.

Because tumor cells under hypoxia have a high demand for AA, we hypothesized that AA transporters may be upregulated selectively to meet this demand. Thus, we investigated the role of HIF-1α regulated AA transporters and in endocrine therapy resistance.

## Results

### Identification of hypoxia-induced amino acid transporters by RNA-sequencing

To define the AA transporters that are involved in hypoxic adaptation, a panel of breast cancer cell lines was cultured in hypoxia (0.1% O_2_) for 24 hours and then RNA-sequencing performed. This panel included estrogen receptor positive (ER+) MCF7, T47D; HER2 positive (HER2+) SKBR3, BT474, ZR751; and triple receptor negative cell lines (TRN) MDA-MB-231, MDA-MB-453, MDA-MB-468, BT20, BT549. We analyzed transmembrane AA transporters expressed in tumors (25, 26). As AAs are crucial for maintenance of the TCA (Krebs) cycle throughout anaplerosis, we applied *in silico* hierarchical clustering and supervised exploratory analysis in order to evaluate which AA transporters, promoting the uptake of specific substrates in the TCA, might be crucial for maintaining anaplerotic pathways in hypoxia. We clustered the AA transporters based on substrate transport: - uptake of glucogenic versus ketogenic AA; essential versus non-essential AA and the entry of their substrates into the TCA cycle.

Our data show that several AA transporters are upregulated under hypoxia in breast cancer cell lines (Supplementary Table 1). Interestingly only three transporters amongst the 25 investigated in breast cancer cells are consistently upregulated and clustered under hypoxia in more than 3 cell lines. *SLC1A1, SLC7A5 (LAT1*) and *SLC38A2* (*SNAT2*) transporters were co-regulated in MCF7, T47D, SKBR3, MDA-MB-231, MDA-MB-468, and BT549 (Figure 1).

**Figure 1.**
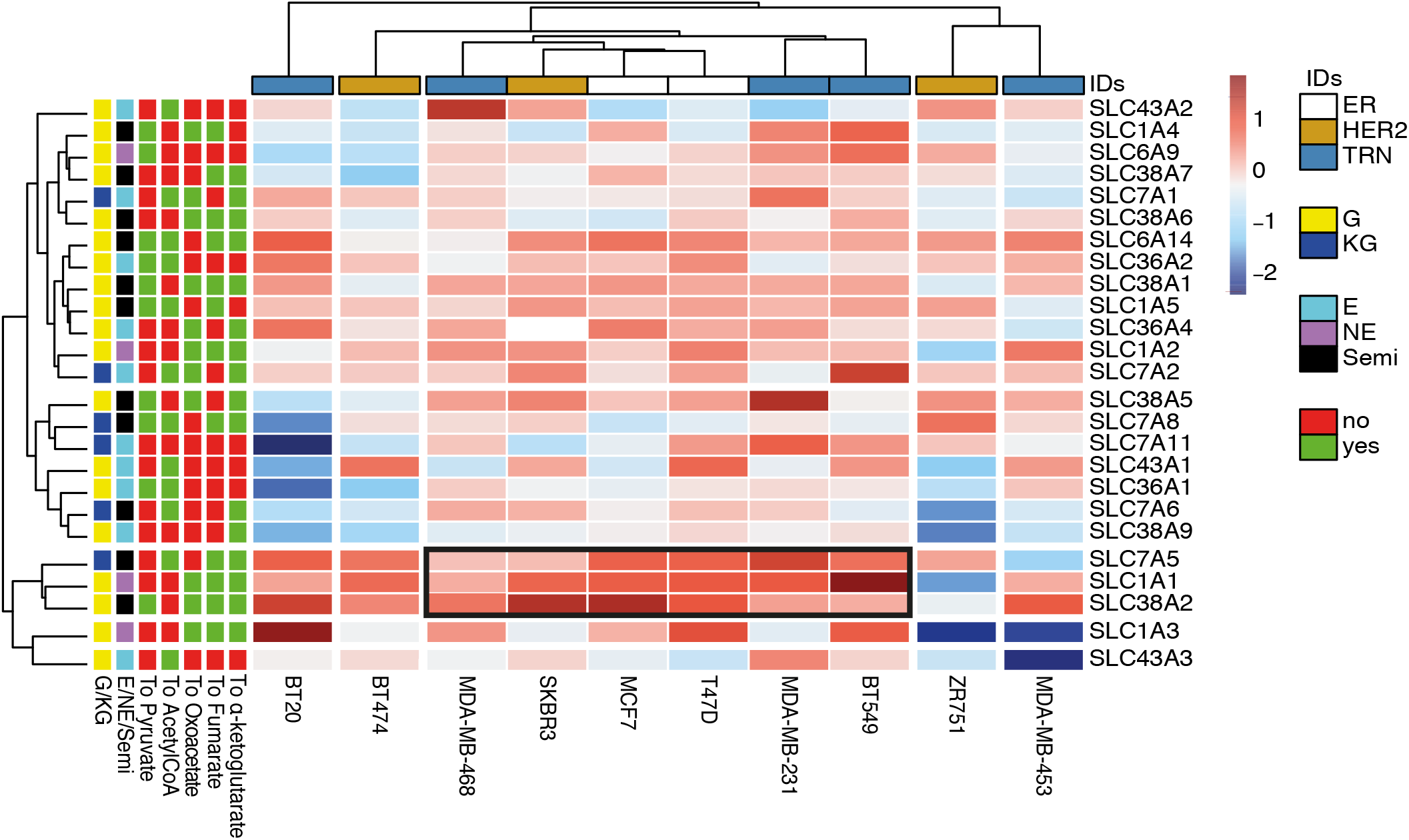
Ten breast cancer cell lines (columns) cultured in normoxia and hypoxia and submitted to RNA-Seq were arranged by supervised average-linkage hierarchical clustering. Heatmap colors represent relative mRNA expression of the selected AA transporters (rows) with higher (red) or lower (blue) expression in hypoxia compared to normoxia. AA transporters that mediate the uptake of gluconeogenic (G) or Ketogenic (KG); essential (E) or non-essential (NE) AAs that enter specific steps of the TCA cycle (α-ketoglutarate, fumarate, oxaloacetate, pyruvate) were clustered. The black box represents the cluster composed of three AA transporters (*SLC1A1, SLC7A5, SLC38A2*) in six cell lines.

The branched-chained AA transporter *LAT1 (SLC7A5*) is a HIF-2α target (27) while *SLC1A1* is a glutamate HIF-dependent transporter (28) (Figure 1). The third AA transporter in the cluster strongly upregulated across a wide spectrum of cell lines under hypoxia is SNAT2, which is involved in neutral AA intake (glutamine, alanine, serine and glycine) (Figure 1) (29). Theoretically the AAs provided by these three transporters could account for all AA precursors of TCA cycle metabolites (Supplemental figure 1A). SLC7A5 can support the uptake of ketogenic AA such as leucine and isoleucine, SLC1A1 the uptake of glutamate and SNAT2 the uptake of other gluconeogenic AA such as glutamine, glycine and serine. Validation of induction by hypoxia by qPCR of these AA transporters was performed in six different cell lines (Supplemental figure 1B) showing the heterogeneity of response to hypoxia.

Further analysis of RNA-sequencing data of four breast cancer cell lines with different receptor phenotypes (MCF-7, SKBR3, MDA-MB-231 and MDA-MB-468) demonstrated that, in addition to the full-length *SNAT2* transcript (isoform 1), there were truncated splice variants of *SNAT2* (isoform 2 and 3) encoding shorter proteins lacking 3 and 6 transmembrane domains (Supplemental figure 1C). The three isoforms encode for proteins (predicted by TMHMM Server v. 2.0) of 56, 45 and 38 kDa, respectively (Supplemental figure 1D). A quantitative PCR assay for specific amplification only of the two main isoforms (1-2) was used, as isoform 3 was not detected. In the cell lines where SNAT2 was induced, there was an increase of expression of both *SNAT2* mRNA isoform 1 and 2 after 24 hours of 0.1% O_2_ (Supplemental figure 1E).

### SNAT2 protein is upregulated under hypoxia in vitro in a HIF-1α dependent manner

We examined SNAT2 protein expression (Figure 2A) using an antibody against SNAT2 isoform 1. Basal RNA (Supplemental figure 2A) and protein and hypoxic SNAT2 induction varied amongst cell lines. Measurement of *SNAT2* mRNA levels in normoxia and hypoxia in a number of additional human cancer cell lines derived from different tissues (colorectal, prostate, ovarian) using qPCR showed that SNAT2 expression is widely upregulated after 48 hours of hypoxia in many cancer cell types (Supplemental figure 2B).

**Figure 2.**
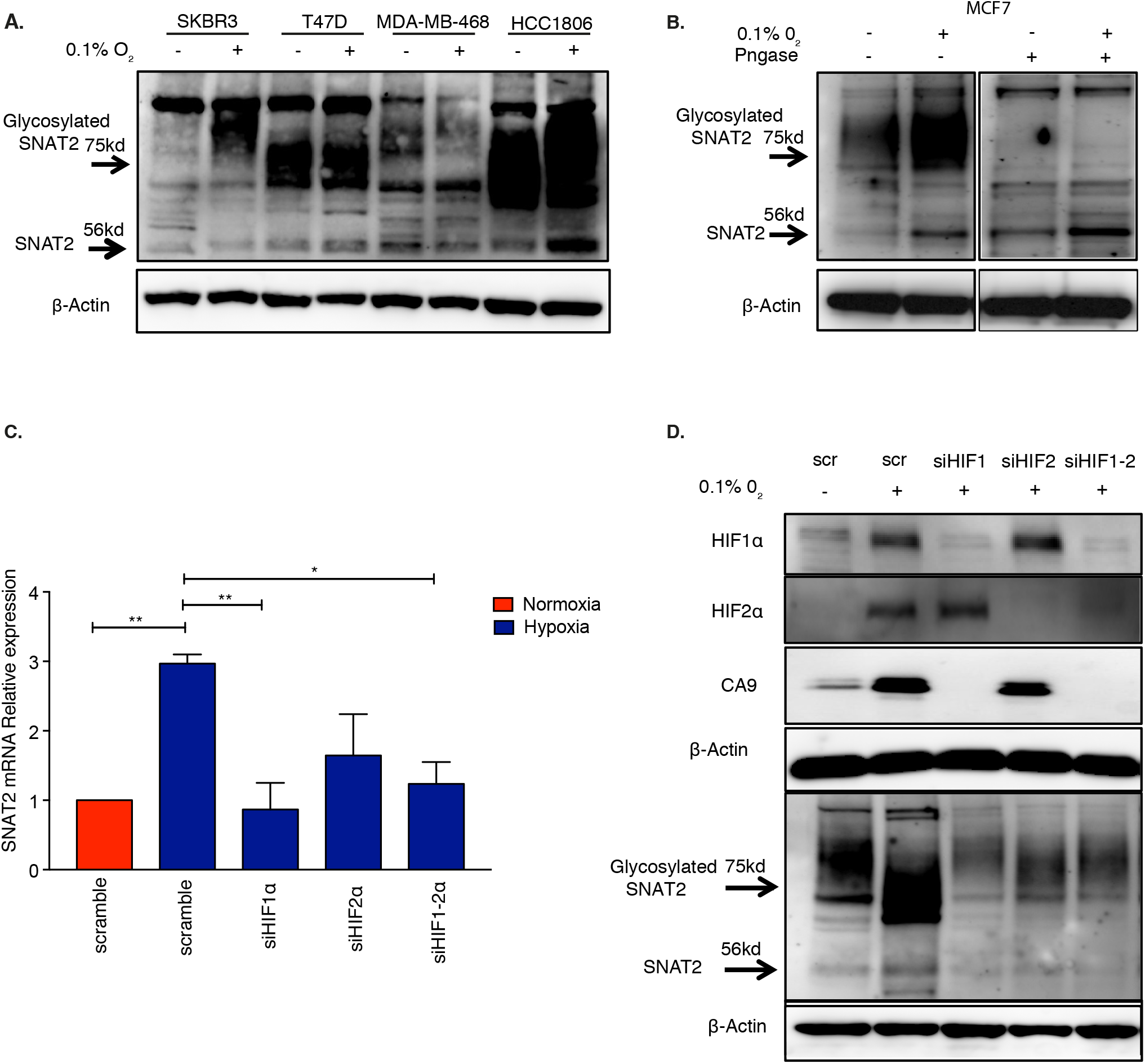
SNAT2 is a hypoxia-induced gene and is regulated by HIF-1α. **(A)** Immunoblot of four breast cancer cell lines from different histotypes showing the heterogeneity in basal SNAT2 expression and their response to 48 hours of hypoxia. **(B)** N-linked deglycosylation assay using PNGase in MCF7 cells lysate in normoxia and hypoxia. **(C-D)** MCF7 cells were transfected with scrambled control (scr), or siRNAs against *HIF-1α* (si*HIF1α*), *HIF-2α* (si*HIF2α*) or both and cultured in normoxia (21% O_2_) or in hypoxia (blue, 0.1% O_2_) for 48 hours. mRNAs were analyzed by RT-qPCR and normalized to 21% O2. *SNAT2* mRNA was normalized to the mean of *β*-actin and RPL11. *p < 0.05 vs. 21% O_2_, **p < 0.01 vs. 21% O_2_. One-way ANOVA test. Error bars, SD. n = 4 **(D)** Immunoblot validation of SNAT2, CA9 (HIF-1α target), HIF-1α and HIF-2α in MCF7 in hypoxia (0.1% O2, 48 hours) after siRNA knockdown specified. β-actin is shown as a loading control

MCF7 ERα positive cells were used for further investigation. In order to determine whether the 75-90 kDa bands detected by western analysis represented a glycosylated isoform of the 56-kDa SNAT2 isoform 1 protein, samples were deglycosylated with an N-linked glycosidase (PNGase F). The disappearance of the 75-kDa bands after PNGase F treatment indicates that this band represents the N-glycosylated form of the molecule (Figure 2B).

The time course of hypoxic induction of SNAT2 in MCF7 cells by qPCR and WB (Supplemental figure 2C-D) peaked at 48 hours, typical of HIF-1α targets. The effect of HIF-1α or HIF-2α depletion by siRNA during this period of hypoxia (Figure 2C-D) was studied. Hypoxic induction of *SNAT2* mRNA expression was significantly reduced by *HIF-1α siRNA* (P < 0.01, n = 3) but not HIF-2α, although a non-significant reduction in SNAT2 induction of ~50% was seen (Figure 2C). The siRNA knockdown was validated by immunoblot of the expression of both HIF-1α and HIF-2α and by measuring carbonic anhydrase 9 (CA9) protein levels, which has been previously shown to be dependent on HIF-1α but not HIF-2α (Figure 2D) (30). To further test the importance of HIF-1α for SNAT2 induction, we stably overexpressed HIF-1α by retroviral vector-mediated transduction in MCF7 (MCF7-HIF1α-o).

Measurement of *SNAT2* and *CA9* mRNA levels in normoxia showed an increased SNAT2 expression in MCF7-HIF1α-o, while no further increase was seen in hypoxia (Supplemental figure 2E).

### SNAT2 is upregulated under hypoxia *in vivo*

To determine whether hypoxic induction of SNAT2 occurs also in solid tumors, two breast cancer cell lines, MCF-7 and MDA-MB-231, were injected into nude mice and grown as xenografts. MDA-231 cells were used to assess effect of hypoxia independently of ER expression. The mice were treated with either saline control or the VEGF inhibitor, Bevacizumab, which slows tumor growth temporarily and induces tumor hypoxia (30). Treatment with Bevacizumab increased SNAT2 protein levels in xenografts derived from both cell lines. In MCF7, SNAT2 protein expression was 10.5% after treatment compared to 6.9% in controls (p<0.05 n=5 per group). In MDA-MB-231, SNAT2 expression was 14.2% after treatment compared to 9.6% in the controls (p<0.05 n=5 per group; Figure 3A). Human *SNAT2* mRNA and *CA9* mRNA were upregulated also (Supplemental figure 3A-B). The levels of CD31 (blood vessels), H/E (necrosis) and CA9 protein (hypoxia) were measured in the same tumors (Supplemental Figure 3C-D). Bevacizumab decreased the percentage of CD31-positive cells. Accordingly, the proportion of necrosis was significantly higher and the percentage of CA9 positive cells was greater in Bevacizumab-treated xenografts compared to PBS (Supplemental figure 3C-D). Furthermore, we found that SNAT2 colocalized in the same area where CA9 was expressed (hypoxic areas) (Fig 3B). To further investigate the effect of HIF-1α on SNAT2 expression in vivo an orthotopic xenograft tumor was established using MCF7-HIF1α-o. Compared with the control, HIF-1α overexpression resulted in an increased SNAT2 expression (Figure 3C-D). These data show that SNAT2 is regulated by hypoxia via HIF-1α *in vivo*, as well as *in vitro*.

**Figure 3.**
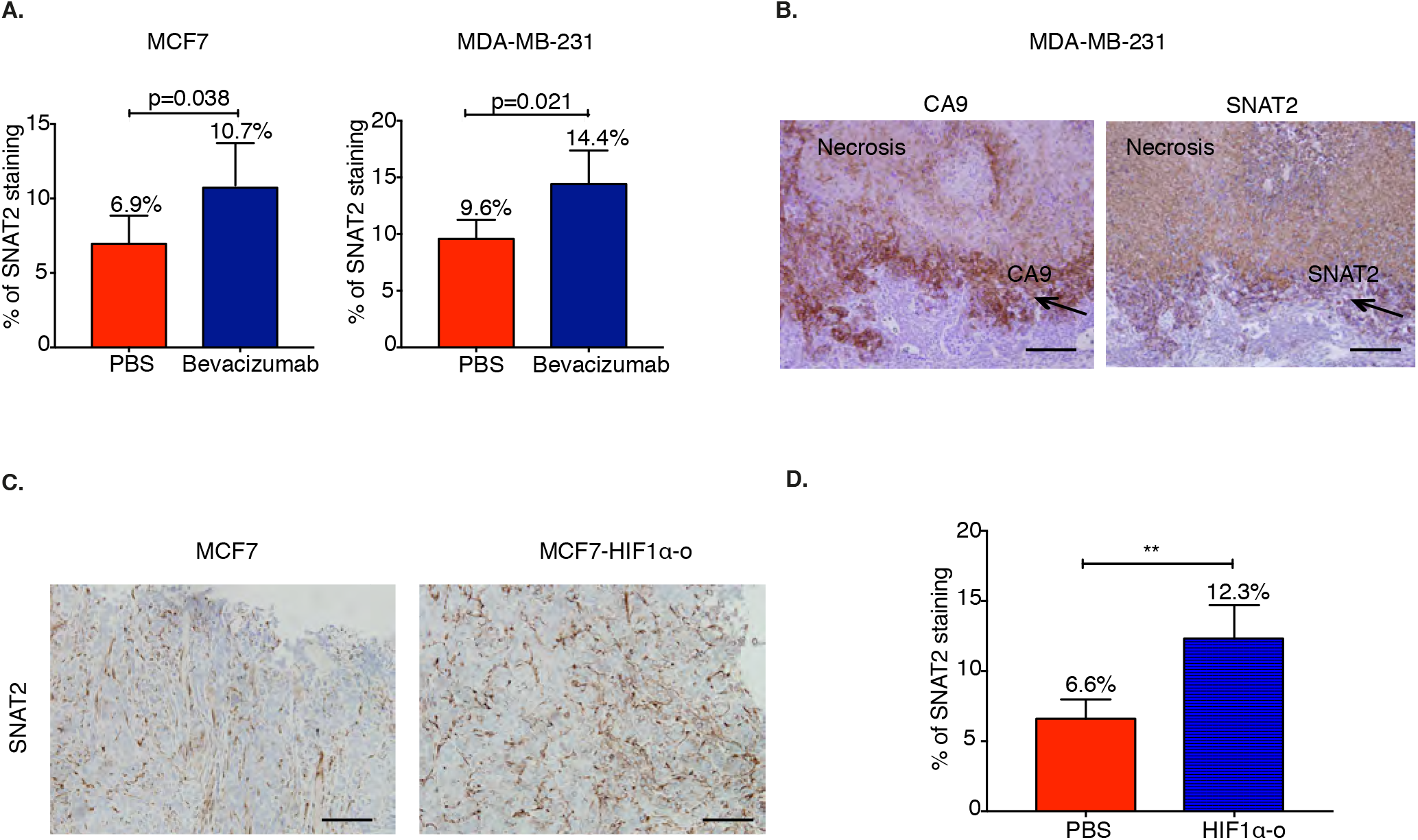
SNAT2 is upregulated *in vivo* by hypoxia and HIF-1α. **(A)** SNAT2 protein expression quantification by ImageJ in MCF7 and MDA-MB-231 xenografts treated with PBS or Bevacizumab. Non-parametric Mann–Whitney test, n=5 PBS and n=5 bevacizumab-treated. **(B)** Representative SNAT2 and CA9 (hypoxia) immunostaining in MDA-MB-231 xenografts showing co-localization of SNAT2 in hypoxic areas of the tumors. Scale bars: 100 μm. (C-D) Representative immunohistochemical images of SNAT2 staining in MCF7 parental and MCF7-HIF-1α-o xenografts and bar chart of scoring. SNAT2 expression was significantly higher in MCF7-HIF-1α-o compared to controls. Scale bars: 100 μm. Non-parametric Mann–Whitney test, n=5 per group. Error bars, SD. ** p < 0.01

### ERα and HIF-1α bind to overlapping sites in the SNAT2 promoter in MCF7 cells, but do not act synergistically

Previous studies showed that the expression of SNAT2 was increased in ER+ positive breast cancer cell lines after 17β-estradiol (E_2_) stimulation and an estrogen response element (ERE) was described in the SNAT2 promoter in rat mammary glands during gestation (31, 32). Because hypoxia is related to adverse outcome in tamoxifen-treated breast cancer patients (14) and SNAT2 was among the 202 genes bound by HIF-1α and ERα and regulated by fulvestrant (14), we investigated the regulation of SNAT2 expression by estrogen and hypoxia in ER+ breast cancer cell lines. Utilizing RNA-seq and HIF-1α and HIF-2α ChIP-seq data in MCF-7 cells, we confirmed the presence of HIF-binding site (HRE) sequences (RCGTG) both for HIF-1α and HIF-2α just upstream of the promoter region of SNAT2 (Figure 4A)(33).

**Figure 4.**
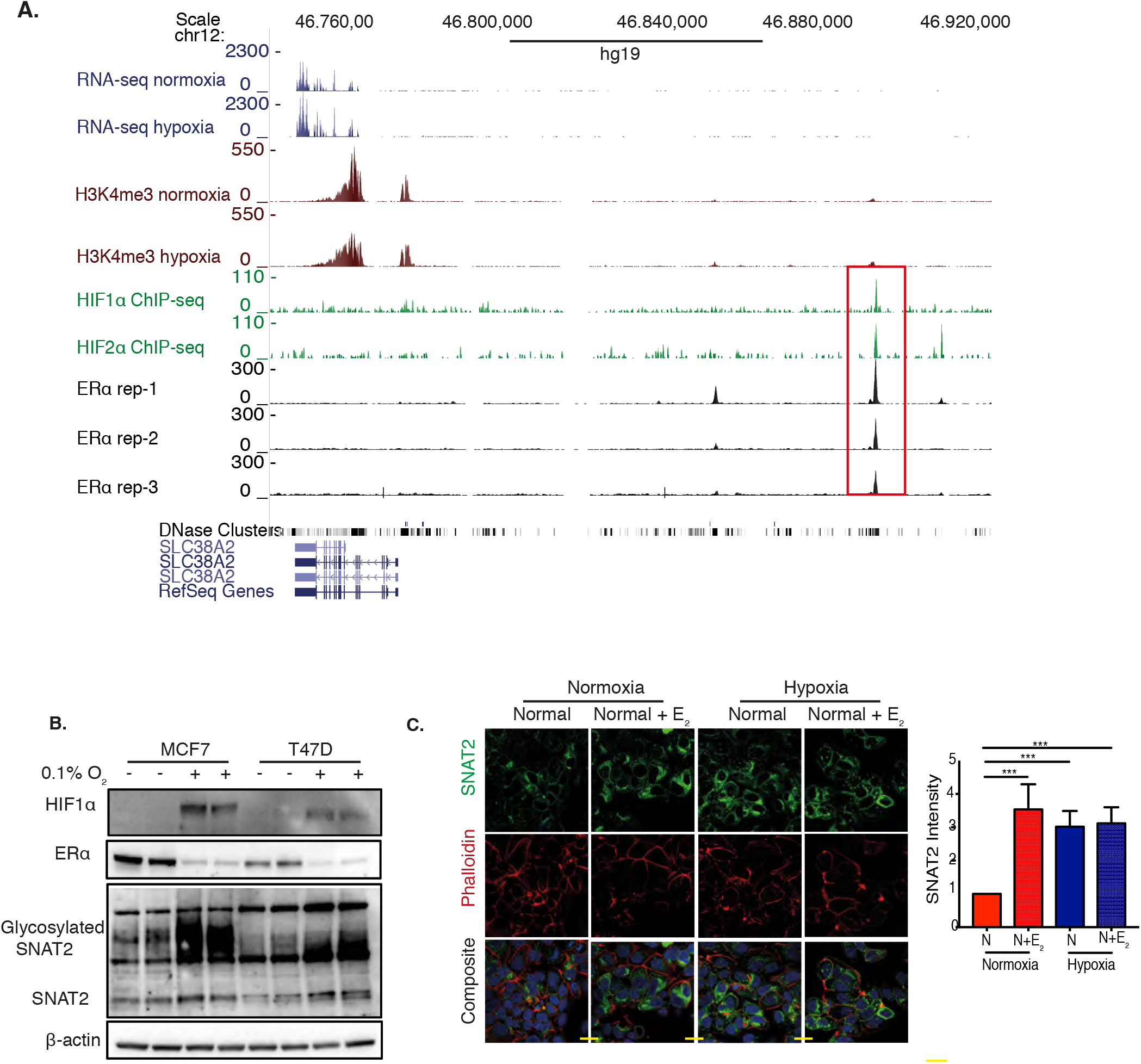
ERα and HIF-1α modulate SNAT2 expression. **(A)** RNA-seq and ChIP-seq genome-browser tracks illustrating occupancy of *HIF-1α* (top), *HIF-2α* (middle) and *ERα* (bottom) at the same genomic coordinates on chromosome 12. The same plots for H3K4me3 are shown. Peaks (red box) represent the areas where transcription factors interact with DNA. **(B)** MCF-7 and T47D cells were treated with 10nM E_2_ in normoxia and hypoxia (0.1% O2) for 48 hours, and cell lysates subjected to western blotting. β-actin is shown as a loading control. **(C)** Confocal microscopy of SNAT2 (green) and phalloidin (F-actin, red) and DAPI (blue) in MCF7 in normoxia and hypoxia (0.1% O_2_) with or without E_2_ treatment (10nM) for 24 hours. Quantification with ImageJ of the staining intensity for SNAT2 in different conditions is presented. Error bars, SD; n = 3.Two-way ANOVA. ***, p < 0.001. *scale bar*, 50 μm.

We aligned the ChIP results of HIF-1α in MCF7 with previous publicly available ERα-ChIP-seq data from ENCODE in the same cell line (34). Both binding sites for HIF-1α and ERα are aligned and overlapping in the genome in cis-regulatory elements, suggesting a potential interaction between these two transcription factors for SNAT2 induction (Figure 4A, Supplemental figure 4A). Interestingly SNAT1, another system A transporter, showed ERα binding sites but not HIFs binding sites (Supplemental figure 4B).

We decided to investigate if an interaction existed between HIF-1α and ERα in driving SNAT2 expression. We cultured MCF7 and T47D cells in normal DMEM medium and then placed them in normoxia or hypoxia and treated with or without E_2_. Estrogen supplementation induced SNAT2 expression in normoxia. Small additive effects were seen in hypoxia in MCF7 but levels of ERα protein significantly decreased (Figure 4B)(35). A similar induction of SNAT2 under estradiol and hypoxia, but with a different magnitude, was also seen for T47D cell line.

Time course experiments of estradiol supplementation in MCF7 in normoxia and hypoxia confirmed the estradiol dependency in normoxia with no significant additive effects in hypoxia at mRNA and protein level (Supplemental figure 4C-D). To evaluate if these different stresses (estradiol and hypoxia) might be responsible for different localization of SNAT2, we examined SNAT2 distribution using immunofluorescence microscopy after 24 hours of hypoxia (0.1% O_2_) and estradiol treatment (10nm). Cells were stained for SNAT2 (green) and the phalloidin (f-actin, red). None of the treatments used significantly affected SNAT2 distribution. SNAT2 was located on punctate structures and the plasma membrane, but the majority was on an intracellular compartment (perinuclear structures), which has been reported to be trans-Golgi network (36, 37). Hypoxia and estradiol resulted in a significant increase in SNAT2 staining intensity but not a clear redistribution of SNAT2 to other structures (Figure 4C).

### ERα and HIF-1α signaling regulate SNAT2 expression independently

To further confirm SNAT2 as an ERα and HIF-1α dependent gene, we cultured MCF7 cells for 5 days in charcoal-stripped phenol-free medium and treated with E2 with or without fulvestrant (ICI182780, which induces ERα degradation) in normoxia and hypoxia. We found that fulvestrant treatment abolished estradiol-dependent SNAT2 induction in normoxia but only a small effect was seen in hypoxia (Figure 5A), suggesting a major role for HIF-1α in this condition.

**Figure 5.**
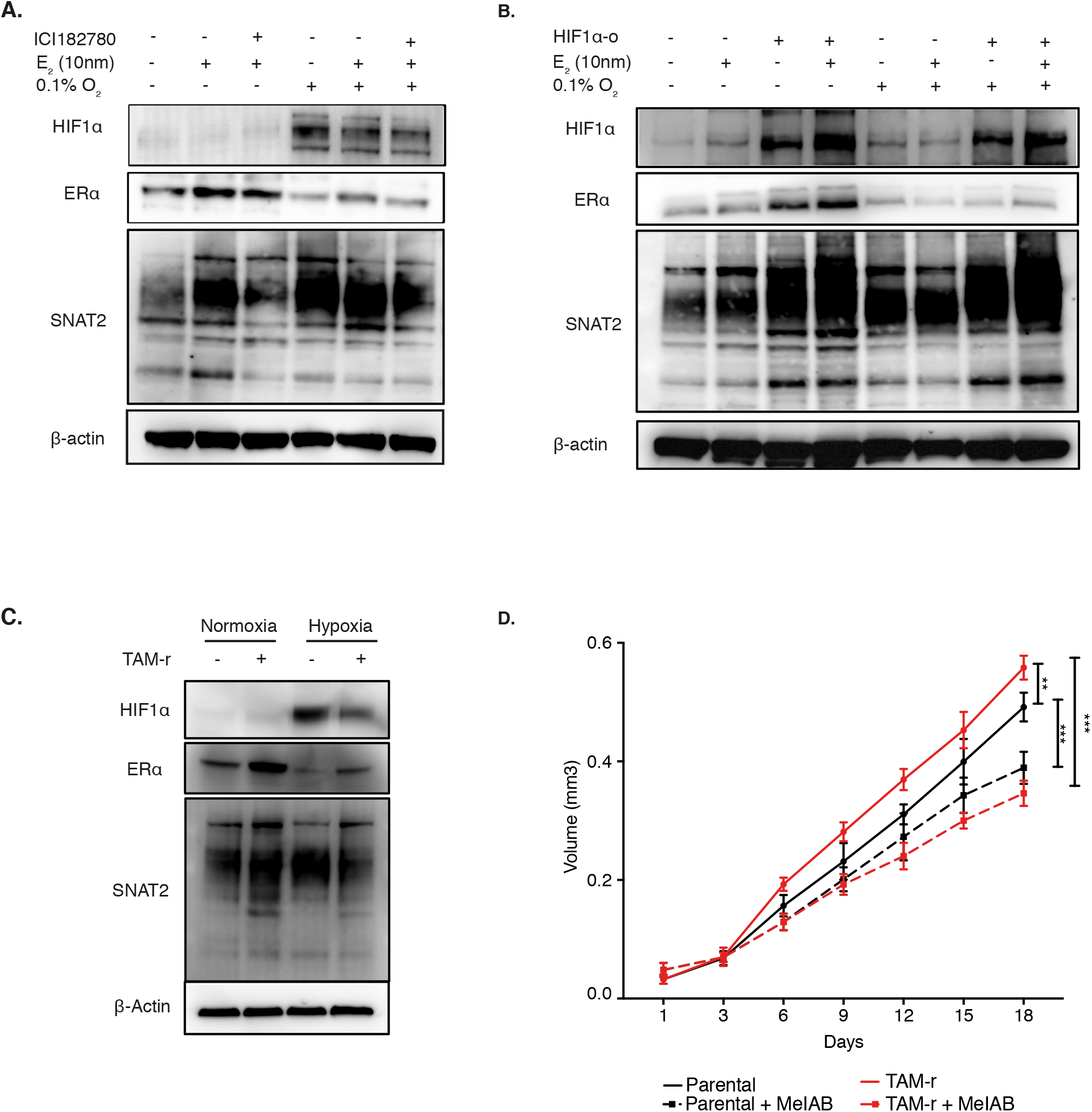
SNAT2 expression is independently modulated by ERα and HIF-1α and is upregulated in a tamoxifen-resistant cell line. **(A)** MCF7 cells were grown in charcoal stripped, phenol-free medium for 3 days and then incubated with or without fulvestrant (ICI 182,780, 10μM) and with or without 10 nM E2 in normoxia and hypoxia (0.1% O2) for 48 hours. β-actin is shown as a loading control **(B)** MCF7 control and MCF7-HIF-1α-o cells were treated with or without 10 nM E2 in normoxia and hypoxia (0.1% O2) for 48 hours. β-actin is shown as a loading control. **(C)** Representative Western blots of parental (MCF7-par) and MCF7 tamoxifen-resistant cells (MCF7-TAMr), which were cultured in normoxia and hypoxia (0.1% O_2_) for 48 hours. β-actin is shown as a loading control. **(D)** Graph of the effect of MeIAB (10 mM) treatment on MCF7 and MCF7-TAMr spheroid growth. Error bars, Two-way ANOVA. SD. **, p < 0.05; ***, p < 0.01; n = 4.

To further test the predominant effect of HIF-1α, MCF7-HIF1α-o cells were treated or not with estradiol in normoxia and hypoxia. SNAT2 was induced in normoxia and hypoxia in MCF7-HIF1α-o cells. When E_2_ was supplemented to WT cells, an increase of SNAT2 was seen only in normoxia but not in hypoxia (Figure 5B). Interestingly when E_2_ was supplemented to MCF7-HIF1α-o an increased level of SNAT2 was seen in both conditions, suggesting that HIF1-o might enhance ER signaling as previously reported (Figure 5B) (14).

To further assess if this overlapping in the genome between HIF-1α and ERα binding sites is only related to SNAT2, we evaluated the 202 genes that we previously showed to have both binding sites for HIF-1α and ERα (14). Using different public available experiments (33, 34), we confirmed 179 genes with both HIF-1α and ERα binding sites. We found only 31 of these genes (31/179, 17.3%) with HIF-1α and ERα binding sites located in the same genomic region (Supplementary Table 1).

Most of these 31 genes, such as ALDOA, NEAT1 and GAPDH are well known HIF-1α or ERα targets.

An example of a gene with overlapping binding sites is QSOX1 (Supplemental figure 5A). To assess if some of these genes had the same pattern of SNAT2 under fulvestrant and hypoxia, we measured mRNA levels in normoxia and hypoxia with or without estradiol and fulvestrant (ICI182780) for *GAPDH, ALDOA* and *NEAT1*. We confirmed also for those genes the estradiol dependency in normoxia and the HIF-1α-dependency in hypoxia and resistance to fulvestrant (Supplemental figure 5B).

This list of 31 genes was selected for further general and metabolic pathway enrichment analysis conducted by MetaCore GeneGo pathway. We found enrichment in glycolysis, HIF-1α and Notch pathways (Supplemental figure 5C). The most relevant GO process was the vesicle-transport pathway (Supplemental figure 5D). These data collectively demonstrate that SNAT2 induction, as well other genes, in MCF7 is regulated by ERα but it becomes predominantly a HIF-1α-dependent gene under hypoxia, due to a potential compensation by HIF-1α when ERα is degraded.

### SNAT2 is increased in a tamoxifen-resistant breast cancer cell line in normoxic conditions

SNAT2 expression was analyzed in the tamoxifen-resistant MCF7 cell line (MCF7/TamR), which has been developed through continuous passage of MCF-7 cells for 6 months in the presence of 4-hydroxytamoxifen (4-OHT) (38). SNAT2 protein was higher in MCF7/TamR compared to parental MCF7 in normoxia, but not in hypoxia (Figure 5C).

When MCF7-TamR cells were grown as a 3D model (spheroids) an increase in cell growth was seen when compared to MCF7 parental. Interestingly when cells were treated with the SNAT2 inhibitor (MeIAB 10mM), MCF7-TamRs were more sensitive to the treatment (maximum volume for MCF7 0.56 mm^3^ vs. 0.35 mm^3^ for MCF7-TamR, two-way ANOVA, p<0.001). This corresponds to a 37.5% decrease in cell growth for MCF7-TamR after MeIAB treatment, versus a 21% decrease in cell growth after MeIAB treatment for parental MCF7 (t-test, p<0.001)(Figure 5D).

### SNAT2 knockdown sensitizes breast cancer cells to anti-estrogen treatment and decreases glutamine consumption, mitochondrial respiration and regulates mTORC1 signaling

To investigate the role of SNAT2 in anti-estrogen resistance, we knocked down *SNAT2* with a siRNA pool (double transfection). Cells were treated with E_2_ (10 nM) and/or fulvestrant (10 μg/ml) in normoxia and hypoxia (Figure 6A). Cell growth was followed over five days. Consistent with their ER positivity MCF7 cells were dependent on E_2_ for their growth in normoxia and sensitive to fulvestrant in normoxia and hypoxia (Figure 6A). SNAT2 knockdown reduced cell proliferation in normoxia but the inhibitory effect was greater in hypoxia (Figure 6A).

**Figure 6.**
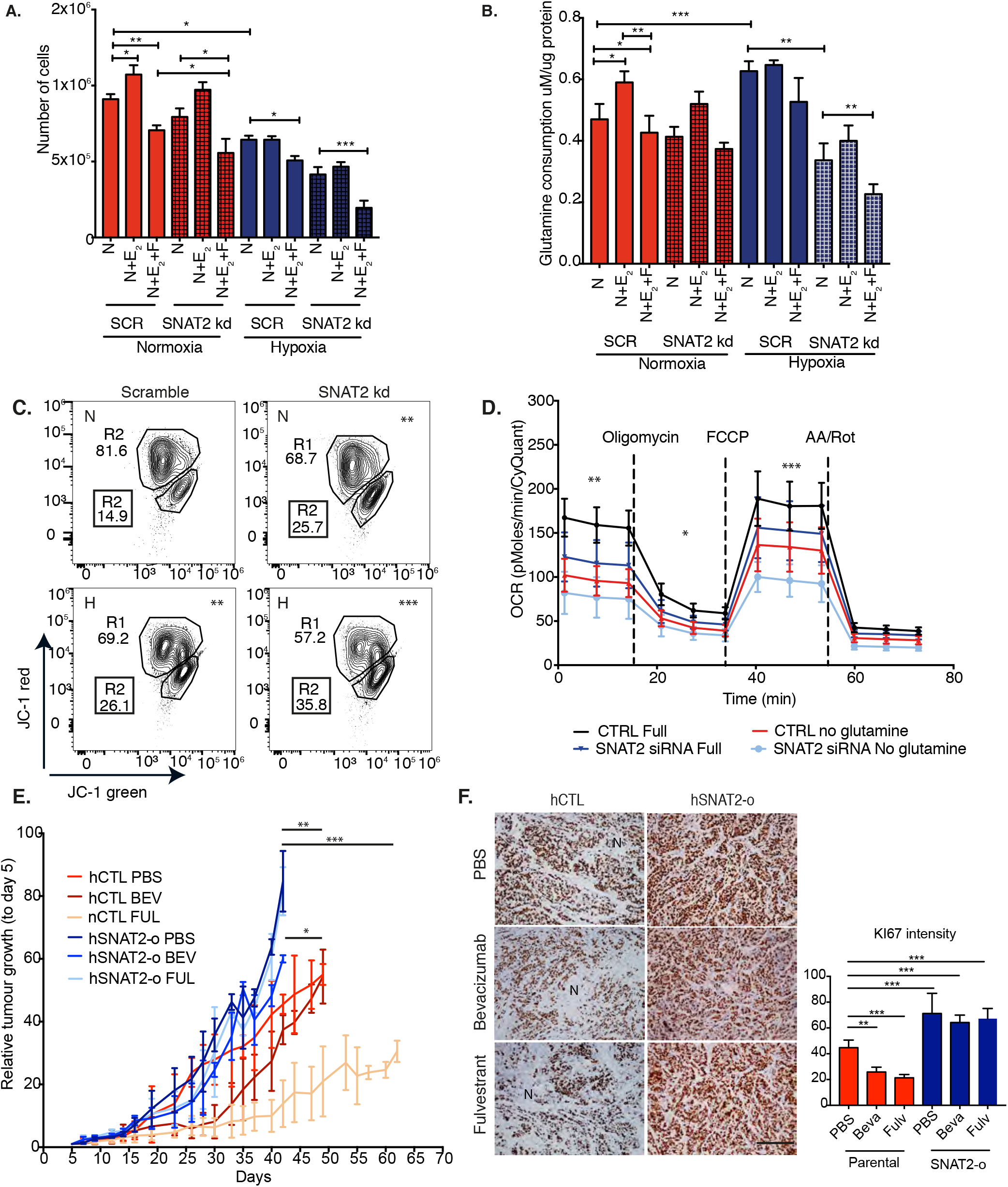
SNAT2 knockdown sensitizes MCF7 cells to fulvestrant treatment in hypoxia and reduces s glutamine intake. **(A)** 10^5^ cells were seeded after 3 days in charcoal-stripped medium and were treated with or without E2 (10 nM), fulvestrant (ICI 182,780, 10μM) and *SNAT2* knockdown for 5 days in charcoal-stripped medium in normoxia and hypoxia (1% O_2_), n = 4. **(B)**. Glutamine consumption was calculated as reported from cells grown in experiments **(A)**. *SNAT2* knockdown reduces glutamine consumption particularly in hypoxia after fulvestrant treatment **(C)**Determination of mitochondrial membrane potential in MCF7 cells by JC-1 staining in normoxia and hypoxia (0.1% O_2_, 48 hours) with or without *SNAT2* knockdown. Values in the trapeziform regions indicate the proportion of cells within them compared to the total number of cells. In the black box (R2) the percentage of depolarized green mitochondria. Chi-square test of different conditions compared to controls **, p < 0.05; ***, p < 0.01; n = 3 **(D)** A representative plot of oxygen consumption measured in MCF7 cells with or without SNAT2 knockdown in normal or glutamine-free medium using the Seahorse Bioscience XF Analyser. Lines indicate time when the oligomycin, the mitochondrial uncoupler FCCP and AA/rotenone were added (E) Xenograft growth curves of MC7 parental and SNAT2-o clones ±Bevacizumab or ±Fulvestrant treatment. Red arrows denote the start of Bevacizumab treatment or Fulvestrant treatment at 150mm3 xenograft volume. Linear regression followed by Student t-test (***p<0.001, **p<0.01, *p<0.05, n=5). Analysis of Ki67 expression by immunohistochemistry with representative images MCF7 parental or SNAT2-o xenografts treated with Bevacizumab or Fulvestrant and a bar chart of the scoring (bottom). Ki67 expression staining is dark brown. N denotes areas of necrosis. Scale bars represent 100μm. One-way ANOVA (***p<0.001, **p<0.01, *p<0.05, n=5).

We then investigated if reduced proliferation was linked to reduced glutamine intake. We found that glutamine consumption was increased after E_2_ supplementation in normoxia and also decreased after fulvestrant treatment as previously described (24). Glutamine consumption was also increased under hypoxia, but no effects were seen with E_2_ supplementation or fulvestrant treatment in hypoxia. Interestingly we found that SNAT2 knockdown did not significantly decrease glutamine consumption in normoxia, but a clear effect was seen under hypoxia. An additive inhibitory effect on reducing glutamine consumption was seen in MCF-7 treated with SNAT2 siRNA (versus scrambled controls) and fulvestrant in hypoxia (Figure 6B). These data indicate that in MCF7, SNAT2 inhibition sensitizes to hypoxia and anti-estrogen treatment and reduces glutamine uptake.

As glutamine is a key metabolite to maintain anaplerosis and mitochondrial function, we used FACS analysis with the cationic dye JC-1 staining to examine the change in mitochondrial membrane potential (ΔΨm) following SNAT2 knockdown in normoxia and hypoxia. Upon knockdown of SNAT2 and hypoxia, the percentage of cells with loss of mitochondrial membrane potential increased from 14.9% of the total to 26.1% of the total, respectively (p<0.01). An additive effect compared to control was seen when SNAT2 knockdown was performed in hypoxia, with 35.8% of the cells showing a loss of mitochondrial membrane potential (p<0.001) (Figure 6D). The effect of SNAT2 knockdown on mitochondria respiration was measured by the SeaHorse analyzer. SNAT2 knockdown reduced basal OCR in normal and glutamine-free medium (Fig 6C). Moreover, SNAT2 knockdown in glutamine-free medium had a more pronounced effect in decreasing basal respiration (p<0.05), ATP production (p<0.01) and maximal respiration (p<0.01) compared to full medium (Figure 6C). Thus, SNAT2 knockdown affects mitochondrial respiration and function in MCF7 breast cancer cells in both normoxic and hypoxic conditions.

We then assessed the SNAT2 knockdown effect on the mTORC1 pathway (a key AA-sensing pathway). SNAT2 depletion both in normoxia and hypoxia reduced the level of phosphorylated mTOR (p-mTOR) its downstream targets, p-pS6 and p-p70 (only in hypoxia) when compared with their total protein levels (Supplemental figure 6A). These results are consistent with previous studies that showed the ability of SNAT2, similar to other AA transporters, to modulate the mTORC1 signaling cascades (39, 40).

Recent research suggested that mTORC1 is localized also at the level of Trans-Golgi network (TGN) and can be activated by specific TGN-AA transporter (41, 42). As SNAT2 is also localized in the TGN (37) and has been suggested to act as a transceptor (40), we performed confocal microscopy of SNAT2, TGN and mTOR in MCF7 cells.

Confocal Immunofluorescence staining of MCF7 cells revealed that SNAT2 is concentrated on the TGN (Supplemental figure 6B). Interestingly, although most mTOR is localized in the cytoplasm, some colocalization with the TGN was also observed, raising the possibility that it might be associated with SNAT2 in this compartment.

### SNAT2 is induced by amino acid deprivation and its overexpression promotes resistance to glutamine starvation, hypoxia and anti-estrogen treatment *in vitro*

To assess if the shortage of specific AAs can upregulate SNAT2, we incubated MCF7 cells in the culture medium depleted of several gluconeogenic and ketogenic AAs in normoxia and hypoxia. Glutamine, serine and glycine (SNAT2 substrates) increased SNAT2 expression in normoxia. Interestingly, when MCF7 cells where incubated under hypoxia, the SNAT2 upregulation became more profound and independent of single AA deprivation (Supplemental figure 6C)

We investigated the growth inhibition after SNAT2 knockdown during metabolic stress (glucose free or glutamine free medium) compared to the nutrient-rich condition in normoxia and hypoxia in MCF7 cell line. SNAT2 knockdown had an effect on 2D cancer cell growth only under hypoxia in full medium (Supplemental figure 6D). SNAT2 knockdown sensitized MCF7 to glutamine deprivation in 2D growth, particularly under hypoxia. A small additive effect was also seen when SNAT2 was knocked-down in glucose-free medium.

We then stably overexpressed *SNAT2* by lentiviral vector-mediated transduction in MCF7 (SNAT2-o) (Supplemental figure 6E). MCF7 SNAT2-0 cells were more resistant to fulvestrant both in normoxia and in hypoxia (Supplemental figure 6F). Reciprocally, MCF7-SNAT2-overexpressing spheroids grew faster than their parental controls in full and, particularly, in glutamine-deprived medium (Supplemental figure 6G). These data indicate that in MCF7, SNAT2 inhibition sensitizes to glutamine deprivation and increased expression enhances growth in low glutamine conditions.

### SNAT2 overexpression promotes resistance to antiangiogenic and antiestrogenic treatment *in vivo*

To further assess the role of SNAT2 in resistance to anti-estrogen treatment and hypoxia, MCF7 and MCF7-SNAT2-o were grown as xenografts with and without bevacizumab or fulvestrant treatment (Figure 6E). MCF7-SNAT2-o xenografts grew faster than the empty vector clone (64.9% the growth rate of the empty vector; *, p < 0.01, n = 7). Bevacizumab treatment decreased the growth of empty vector tumors in a first phase and then tumor became resistant. Fulvestrant treatment also reduced xenograft growth rate (55.4% the rate of empty vector control; *, p < 0.01, n = 7). The SNAT2 expressors grew more rapidly than the controls and there was no impact on this increased growth by either treatment.

The MCF7-SNAT2-o clones treated with bevacizumab grew significantly faster than MCF7 empty vector treated with bevacizumab (11.2% the growth rate of treated empty vector; *, p = 0.038, n =7). More strikingly MCF7-SNAT2-o clones treated with fulvestrant grew significantly faster than MCF7 empty vector and became completely resistant to fulvestrant (267% the growth rate of treated empty vector, p < 0.001, n =7) with a curve similar to untreated MCF7-SNAT2-o.

SNAT2-o xenografts, independently of the treatments, had increased proliferation compared to controls as determined by Ki67 staining (p<0.001). Interestingly no significant differences were seen in Ki67 staining amongst SNAT2-o xenografts treated or not with bevacizumab or fulvestrant (Supplemental figure 6F).

### SNAT2 expression correlates with tumor hypoxia and is associated with a poorer recurrence free survival in endocrine treated breast cancer patients

To evaluate the clinical relevance of SNAT2 in breast cancer, we first assessed if SNAT2 expression correlates with hypoxia *in vivo*. Analyzing gene expression data from 2433 breast cancer patients using the Metabric cohort (43), we found that *SNAT2* mRNA abundance significantly correlated with the expression of many genes in our previously reported *in vivo* hypoxia signature, (44) but not with the whole signature itself (Supplemental figure 7A). For confirmation *SLC7A5* (a HIF-2α regulated gene) showed a correlation with the hypoxia signature (Supplemental figure 7A). Interestingly *SNAT2* mRNA levels did not correlate with *c-Myc* copy number, a key regulator of glutamine metabolism (Supplemental figure 7B).

We used additional expression data derived from different breast cancer gene-array cohorts (4132 patients)(45) and we tested if the *SNAT2* levels correlated with worse outcomes in ER+ patients treated with all adjuvant endocrine treatments [including aromatase inhibitors] or tamoxifen only. High *SNAT2* expression correlated with low recurrence free-survival in patients who received endocrine or tamoxifen treatments (Figure 7A). Moreover, when we looked at the tamoxifen-treated cohort we found that high *SNAT2* levels correlated with worse outcome in tamoxifen-treated luminal B but not luminal A patients (Figure 7B).

**Figure 7.**
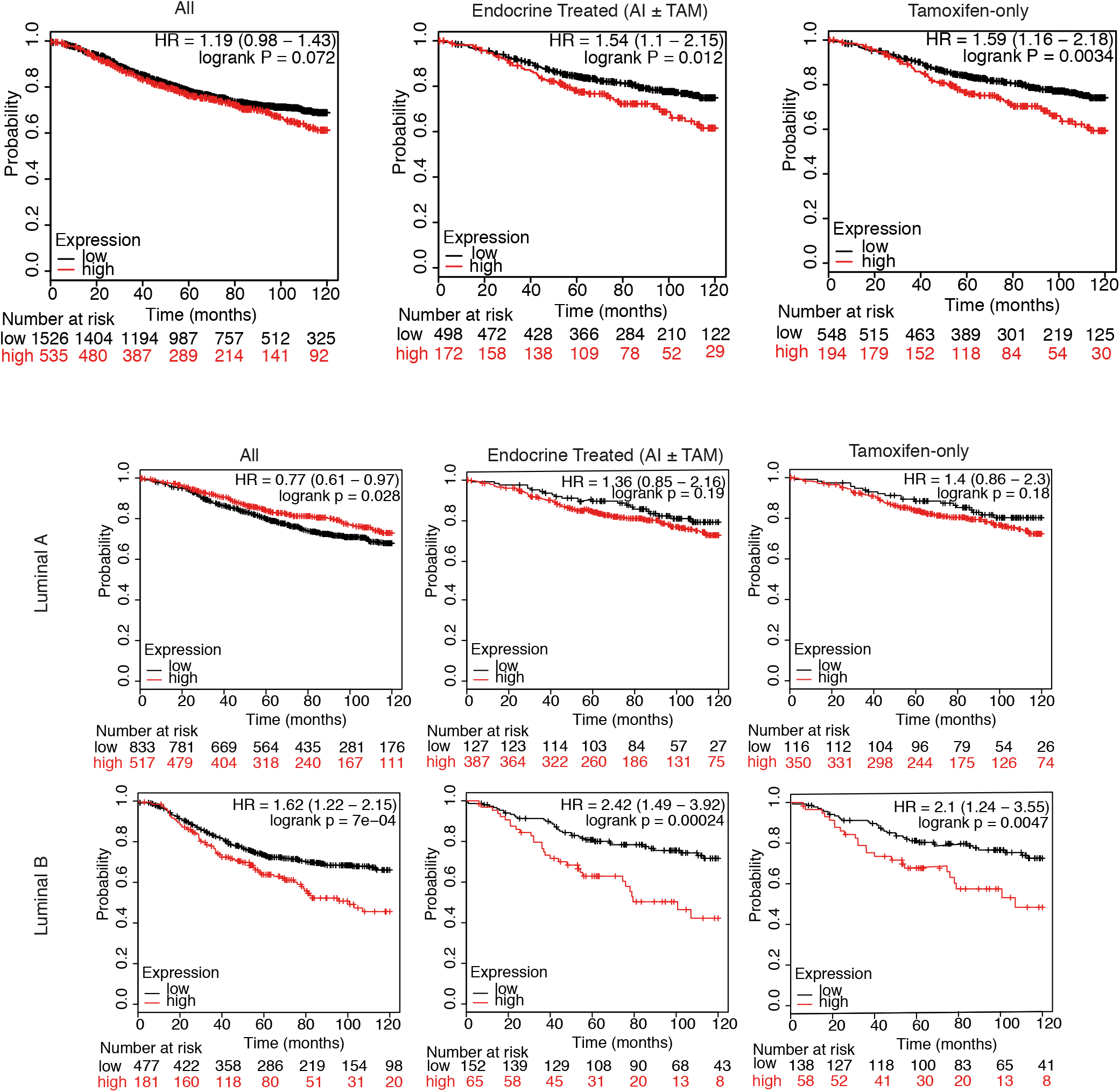
SNAT2 expression is related to poor outcomes in endocrine-treated breast cancer patients. **(A)**Kaplan–Meier plot for ER+ breast cancer patients from different gene-array studies stratified according to expression of *SNAT2* mRNA (above [red] versus below [black] third quartile) (left panel). High tumor SNAT2 levels are associated with decreased recurrence-free survival only in patients who received endocrine [aromatase inhibitor (AI) or tamoxifen (TAM), middle panel] or tamoxifen (right panel) as part of their treatment for breast cancer. **(B)** Kaplan–Meier plot for luminal A and B breast cancer patients (left panel), and for same subgroups who received endocrine [aromatase inhibitor (AI) or tamoxifen (TAM), middle panel] or tamoxifen (right panel) as part of their treatment for cancer. Patients were stratified according to expression of *SNAT2* mRNA (above [red] versus below [black] third quartile). Recurrence free survival was evaluated

## Discussion

Hypoxia is a cause of tumor aggressiveness and resistance to treatments, including endocrine therapy. Although the role of hypoxia and the HIF-1α transcriptional response in promoting tumor progression and metastasis is well established, the direct contribution of the HIF family to the regulation of AA transporters has been less studied, apart from LAT1 and glutamate transporters (25). Here we describe a new mechanism by which hypoxia can produce resistance to anti-endocrine therapy, by substituting for ERα in regulating the AA transporter SNAT2, a glutamine transporter. We found that despite the induction of several AA transporters in hypoxia, SNAT2 expression was key and able to control the growth response to glutamine and anti-estrogen treatment *in vitro* and markedly so *in vivo*.

Both main SNAT2 isoforms were hypoxia-regulated. Until now the main isoform has been investigated while for other isoforms their role has not been studied yet. Recently, it has been shown that different SLC1A5 isoforms are upregulated and needed for SLC1A5 activity under AA deprivation, suggesting that the isoforms might form a complex necessary for AA transporter activity (46). Therefore, the role of isoform 2 now requires further investigation. The maturation of SNAT2 protein and transmembrane localization requires its glycosylation (47) which was maintained in hypoxia.

Accumulating data suggest a significant interplay between hypoxia and estrogen-mediated pathways in breast cancer cells (14, 48, 49). Hypoxia down-regulates ERα in human breast cancer cells via a proteasome pathway (49). We found that both HIF-1α and ERα have binding sites on the same cis-regulatory region of the SNAT2 gene, suggesting that SNAT2 transcription can be regulated by either of these two transcription factors, permitting hypoxia-dependent growth under anti-estrogen treatment. Of interest was a further set of genes regulated by ERα in normoxia and by HIF-1α in hypoxia independently of ERα. These genes represent several pathways with a well-recognized role in tumor growth and warrant further investigation for roles in hormone independence.

Several observations support the role of SNAT2 in hormone resistance *in vitro*. Depletion of SNAT2 increased the inhibitory effects of fulvestrant and its overexpression decreased the effects. The tamoxifen-resistant MCF7-TAM cell line had an increased SNAT2 expression. Pharmacological inhibition of SNAT2 had a greater effect on 3D spheroid growth of MCF7-TAM compared to parental MCF7.

The xenograft models demonstrated a far more profound effect than the *in vitro* data. It is well known that metabolic modifications are often more profound *in vivo* because of continuous poor nutrient conditions and hypoxic, acidic and glucose gradients. Additionally, SNAT2 expression is tightly controlled also at the post-translational level (50). AA starvation results in uncharged tRNAs, activating GCN2 and ATF4, which in turn will cause translation of the abundant SNAT2 mRNA by a cap-independent mechanism (51). SNAT2 protein is also degraded after ubiquitination (52), and this may occur under nutrient-replete conditions, but is halted when amino acids are depleted. This could explain the more rapid growth rate of SNAT2 overexpressing tumors and potentially related to scavenging glutamine.

SNAT2 knockdown shifted the mitochondrial membrane potential both in normoxia and hypoxia, suggesting the decrease in cell growth might be mediated by mitochondrial impairment. These data suggest that the dependency on SNAT2 in hypoxia to maintain the TCA cycle cannot be compensated by other AA transporters (such SNAT1 or ASCT2), which are not induced by hypoxia in breast cancer cells.

c-Myc is a well-known downstream effector of ER-α (53). It is upregulated by E_2_ and it plays a critical role in modulating resistance to endocrine therapies, by the unfolded protein response (UPR), in ERα-positive breast cancer (54-56). Although SNAT2 is induced by the UPR (57) and Myc was found to selectively bind to the promoter regions of SNAT2 (58), we did not find any correlation between *c-Myc* copy number and *SNAT2* mRNA levels in breast cancer patients (Supplemental figure 7B).

Finally, our analysis of clinical data available from large RNA expression cohorts suggests that high baseline SNAT2 expression correlates with reduced recurrence free survival in ER+ endocrine-treated breast cancer patients, specifically in luminal B subtype. This is particularly interesting as the luminal B subtype (ER+ subtype) has aggressive clinical behavior, with prognosis similar to that of HER2-enriched and basal-like group and a lower sensitivity to endocrine treatment compared to luminal A ER+ breast cancer (59). Interestingly, recent studies showed that a high expression of different AA transporters, such as SLC3A2 or SLC7A5, is related to poor outcomes in luminal B ER+ breast cancer subtype (60, 61). These data suggest that this subtype might be particular vulnerable to glutamine depletion.

Clinical selection may be possible, as recently an amino acid-based PET radiotracer, ^18^F-fluciclovine, has been investigated in the imaging of breast cancer (62).

As targeting glutamine metabolism is the subject of intensive research due to its potential clinical applications (20), we propose that SNAT2 should be investigated as a predictive biomarker and a potential target for well-defined molecular subtypes (luminal B) of ER+ breast cancer patients and as a combination approach to overcome endocrine-therapy resistance.

## Disclosure of Potential Conflicts of Interest

No potential conflicts of interest were disclosed by the authors.

## Authors’ Contributions

ALH and MM conceived and designed the study. MM, HC, DJ, ET conducted the experiments described in the paper. HS helped with the cloning. EB, MM, SL, and AM performed the xenografts experiments. EB was responsible for immunohistochemistry. FB, WCC, SH, NG and AV performed the bioinformatics analysis and clustering. DCIG, AV and ALH, provided critical insights and/or conceived and designed the experiments. MM and ALH wrote the manuscript. All authors reviewed the manuscript and agreed with results. ALH supervised and coordinated the work.

## Acknowledgments

We thank Dr. Ioanna Rota for FlowJo analysis and her help in the revision of the manuscript. Dr. Christoffer Lagerholm for help with confocal microscopy. This research was supported by funding from Cancer Research UK (ALH), Oxford NIHR Biomedical Research Centre (ALH), Breast Cancer Research Foundation (ALH), Friends of Kennington Cancer Fund (ALH), Oxfordshire Community foundation (M.M) and by a Fondazione Veronesi fellowship (to M.M.).

This study makes use of data generated by the Molecular Taxonomy of Breast Cancer International Consortium, which was funded by Cancer Research UK and the British Columbia Cancer Agency Branch.

## Materials and Methods

### Cell culture

Cell lines were available from ATCC (MCF7, T47D, SKBR3, BT474, HCC1806, MDA-MB-231, SUM159, MDA-MB-468, PC3, HCC1954, PC3, DU159), Clare Hall Laboratories (HCT116) or kindly gift of the Prof Ahmed Ahmed laboratory (SKOV3, HEY, OC316). Cell line authentication was carried out by STR analyses (LGC Standards) 6 months prior to the first submission of the manuscript. Cells were maintained in a humidified incubator at 5% CO2 and 37°C. For hypoxic exposure, cells were grown in a humidified atmosphere of 0.1% or 1% O2, 5% CO_2_ at 37°C. All cell lines were maintained in DMEMα 25mM glucose supplemented with 10% FBS. All the experiments were performed in 5mM glucose. For spheroid culture, aggregation was initiated by plating 1,000–5,000 cells into ultra-low–adherent round-bottom 96-well plates (VWR) with (MCF7) or without Matrigel (HCT116) and centrifuging these at 2,000 × g for 5 minutes.

### RNA sequencing and bioinformatics

RNA-sequencing was performed as previously described (33). The sequenced paired-end reads were aligned to human reference genome GRCh38 with transcriptomic information of the genome by Bowtie 2.2.6 and Tophat v2.1. We then estimated the fold changes on the normalized expression level FPKM (Fragments Per Kilobase of transcript per Million mapped reads) for each gene and its transcriptional isoforms from those mapped reads using Cuffdiff 2.2.1. The means and standard deviations are calculated over the triplicates of each cell line. ChIP-seq databases were analyzed as previously described (63). For heatmap and hierarchical clustering, supervised exploratory analysis, transcriptomic data were standardized by z-score scale. K-mean was utilized to determine the optimal number of clusters. Survival analysis of Metabric cohort was performed in R statistical environment (v3.3.2) using survival package (v2.41-3).

### Gene silencing by RNA interference

Reverse transfection of siRNA duplexes (20□nm) was in Optimem (Invitrogen, Waltham, MA, USA) using Lipofectamine RNAiMax (Invitrogen, Carlsbad, CA, USA) according to the manufacturer’s instructions. The oligonucleotides are reported in the Supplementary material.

### RNA extraction and RT-PCR

Cells were lysed in Trizol reagent (Invitrogen), and RNA was extracted using ethanol precipitation. RNA quality and quantity were confirmed using the NanoDrop ND-1000 spectrophotometer (Wilmington, USA). Reverse transcription (RT) was performed using the High Capacity cDNA RT kit (Applied Biosystems, Life Technologies, Bleiswijk, Netherlands). QrtPCR was performed using the SensiFAST SYBR No-ROX Kit (Bioline, London, UK). Raw data were analyzed using Ribosomal Protein L11 (RPL11) and β-actin as a housekeeping gene. Each PCR reaction was done in triplicate. Primer sequences are reported in Supplementary Material.

### Cell Proliferation

Approximately 100,000 cells per well (MCF7 and T47D) were seeded in triplicate on 6-well tissue culture plates. Each triplicate was treated with 10 μM Fulvestrant (Sigma-Aldrich, I4409-25MG) or treated with SNAT2 siRNA (20nM) and maintained under 21% or 1% O_2_ for 5 days. Trypsinized cells were counted using Cellometer Auto T4 Cell Counter (Nexcelom Bioscience). Only the cell population with the size of 13–27 μm was counted. We used α-(Methylamino) isobutyric acid (MeIAB) (M2383, Sigma-Aldrich) at 10mM concentration in MCF7-TAM spheroids. Treatment was started at day 3 and renewed every two days. Pictures were taken every 2-3 days.

### Generation of SNAT2 and HIF-1α overexpressing MCF7

Stable MCF7 control or overexpressing the full-length SNAT2 cell lines were generated using viral infections. Briefly, 30 μg of empty vector-containing pLenti6.2-V5 plasmid or full length SNAT2 cDNA-containing pLenti6.2-V5 plasmid with 15-μg pPAX2 and 5μg of pMDG.2 (Addgene) were transfected into HEK293t packaging cell line using the CaCl_2_ method. The viral supernatant was recovered and the transduced cells were generated by infection at 5 MOI (multiplicity of infectious units) and selected with 1 μg/mL of Puromycin for 1 week. SNAT2 expression was confirmed by Western blot. The generation of MCF7 HIF-1α overexpressing cells was previously reported(14)

### Immunoblotting

Protein concentrations were quantified using a BCA protein assay kit (Thermo Fisher Scientific c, Cramlington, UK). Samples containing 20μg of protein were in Laemmli buffer and separated using 8-12% pre-cast SDS-PAGE gels (Bio-Rad, USA). Proteins were transferred to PVDF membranes (Millipore, USA), blocked for 1 hour in transfer buffer (25 mM Tris, pH 7.5, 0.15 M NaCl, 0.05% Tween 20) 5% milk. Primary antibodies were used at 1:1.000 unless otherwise stated. Rabbit anti-SNAT2 (BMP081, MBL International), mouse anti-HIF1α (BD Biosciences, Clone 541, 610958, 1:500), mouse anti-CA9 (M75, Ab00414-1.1, Absolute); rabbit anti-HIF2α (NB 100-122, Novus Biologicals), rabbit anti-ERα (D8H8, 8644, Cell Signalling) and actin-HRP (Sigma). SNAT2 was deglycosylated with PNGase F (New England Biolabs, Hitchin, UK; no. P0704) according to the manufacturer’s instructions. Appropriate secondary horseradish peroxidase–linked antibodies were used (Dako). Blots were developed on Kodak lm (Sigma, Steinheim, Germany) using ECL plus reagent (GE Healthcare, UK). Image J software was used to quantify bands intensities after β-actin normalization.

### Immunofluorescence

Immunostaining were performed as previously reported (64). Samples were incubated overnight with SNAT2 primary antibodies and then incubated with fluorescently conjugated donkey secondary antibodies and DAPI for 1 h at 37°C followed by a further three 10-min washes in PBST. All samples were imaged on a Zeiss LSM 780 confocal microscope. Magnification images were obtained using the x40 objective (1.4NA, Oil DIC, Plan-APOCHROMAT). Image J software was used to quantify SNAT2 intensity as previously described (65).

### Immunohistochemistry

Immunohistochemistry was carried out as previously described (66). The following primary antibodies were used at overnight incubation: Pimonidazole (Hypoxyprobe-1; Chemicon International, 1:100), anti-mouse CD31/PECAM (1:100, Clone ER-MP12, ThermoFisher) mouse anti-CA9 (M75, Absolute antibody, 1:100), rabbit anti-SNAT2 (BMP081, MBL International, 1:100). A standard haematoxylin and eosin protocol was followed to assess the morphology of HCT116 spheroids and the amount of necrosis on xenografts. Slides were incubated with the antimouse secondary antibody (Dako) for 45 minutes and washed in PBS. 3,3'-Diaminobenzidine (DAB, Dako) was applied to the sections for 7 minutes. The slides were counterstained by immersing in haematoxylin solution (Sigma-Aldrich) and mounted with Aquamount (VWR). Secondary-only control staining was performed routinely. Necrosis was quantified histologically on haematoxylin-stained sections as previously described (66). Expression of CA9 and necrosis was quantified on whole sections by using ImageJ software (US National Institutes of Health, Bethesda, MD, USA). Expression of CD31 was quantified by using 15 random fields of slides at × 100 magnification.

### Determination of glutamine consumption

Cells (1×104/well) in a 24 well plate were cultured for 24 hours in medium without phenol red, medium was collected, and cells were lysed with RIPA buffer (Sigma-Aldrich). Concentrations of glutamine in the medium and in the cell lysate were determined with the Glutamine Detection Assay Kit (Abcam, ab197011). A standard curve was determined for each experiment to calculate the concentration of glutamine in samples as per manufacture guideline. Glutamine levels were calculated and normalized to total protein levels. The glutamine level of normal culture medium was also measured, and the glutamine consumption was calculated as (glutamine in normal medium-glutamine in medium after culturing cells) and normalized to protein level.

### Seahorse XF-24 metabolic flux analysis

Oxygen consumption rates and extracellular acidification rate were measured at 37 °C using an XF24 extracellular analyzer (Seahorse Bioscience). MCF7 were plated in 24-well plates for 24 h (50,000 cells per well) in DMEM 5mM glucose, 4mM glutamine with or without SNAT2 knockdown. 24 hours before the experiments cells were changed to normal 5mM medium or medium with no glutamine. Mitochondrial function was interrogated by the sequential injection of oligomycin A (1.5 μM, which inhibits ATP synthesis), FCCP (0.5 μM, uncoupling agent) and antimycin A (10 μM, complex III inhibitor) in combination with rotenone (2 μM, complex I inhibitor), as per protocol. This allowed for the calculation of ATP-linked O2 consumption, proton leak, maximal respiratory capacity, reserve capacity and non-mt respiration. After each experiment, the protein concentrations in each well were measured by Bio-Rad DC™ Protein Assay. Values were normalized to protein concentration after the completion of the XF assay. Three baseline measurements were taken before sequential injection of mitochondrial inhibitors. Oxygen consumption rate was automatically calculated by Seahorse XF-24 software. Every point represents an average of n = 6.

### Mitochondrial Permeability Potential

Cells were stained with the cationic dye JC-1 (Thermofisher), which exhibits potential-dependent accumulation in mitochondria. At low membrane potentials, JC-1 continues to exist as a monomer and produces a green fluorescence (emission at 527 nm). At high membrane potentials or concentrations, JC-1 forms J aggregates (emission at 590 nm) and produces a red fluorescence. Briefly, cells were grown as indicated and incubated in 1.25 □ μm JC1 and analyzed by the LSR-II (BD) flow cytometer. The gating strategy was based on the presence of three cell populations with high (R1) versus low (R2) emission in the PE channel. Data were processed by FlowJo v7.6.4.

### Xenograft studies

Procedures were carried out under a Home Office license. Xenograft experiments were performed in female (1) BALB/c nunu (MCF-7 cells), (2) BALB/c SCID (MDA-MB-231). A total of 2.5 × 10^6^ (MCF-7) cells or 10 × 10^6^ (MDA-MB-231 cells were injected subcutaneously in the lower flank in equal volumes of Matrigel (BD Biosciences, Oxford, UK). Mice injected with MCF-7 cells had estrogen (5□μg/ml) added to their drinking water. Once tumors reached 150□mm^3^, mice received either intraperitoneal (i.p) bevacizumab (10 □mg/kg every 3 days) or vehicle control for (1) and (2) experiments. For MCF7 ± HIF-1 xenografts, 50 μL of Matrigel containing 5 × 10^6^ MCF7 ± HIF-1α were implanted into the mammary fat pad of NSG female mice, aged 5–6 wks, implanted with estrogen (0.72 mg; 90-d release; Innovative Research of America). When tumors reached 1.44 cm^3^, mice were sacrificed by cervical dislocation. (3) For MCF7 parental and MCF7 SNAT2-o experiments 5 × 10^6^ cells were injected subcutaneously in the lower flank in equal volumes of Matrigel. Bevacizumab was injected i.p. every 3 days (10mg/kg) and fulvestrant (), until sacrifice. All xenografts were also implanted with estrogen as seen above. Tumor growth was monitored three times per week measuring the length (L), width (W) and height (H) of each tumor using calipers. Volumes were calculated from the formula 1/6 × π × L × W × H.

### Bioinformatics and statistical analysis

For heatmap and hierarchical clustering, unsupervised exploratory analysis RNA transcriptomic data k-mean was utilized to determine the optimal number of clusters.

Statistical analysis and graphs were performed using GraphPad Prism v6.0 (GraphPad, La Jolla, CA, USA). Results are plotted as mean values with standard deviation (SD). Statistical tests and the number of repeats are described in the figure legends. Student’s t-test was used for two sample analyses and normal distributions were assumed, otherwise the non-parametric Mann–Whitney test was used. Analysis of variance was used for >2 sample analyses. No samples or experimental repeats were excluded from analyses. No statistical methods were used for the samples size selection of other experiments

---

**Supplementary figure 1. SNAT2 is increased by hypoxia in breast cancer cell lines (A)** Schematic model showing that the three clustered AA transporters (red) upregulated under hypoxia can potentially fulfill all the AA precursors of TCA cycle metabolites. These three AA transporters can provide ketogenic AA (SLC7A5), gluconeogenic AA (SLC38A2) and glutamate and aspartate (SLC1A1) intake. These AA can enter the TCA cycle at the level of all the TCA cycle intermediates (Pyruvate; Acetyl-coA; Citrate; α-ketoglutarate; Succinyl-CoA; Fumarate; OAA, oxaloacetate) throughout reactions of decarboxylation and transamination. **(B)** The relative expression of different AA transporters in hypoxia (0.1% O_2_, 48 hours; blue bars) compared to normoxia (red bars) in 6 breast cancer cell lines. **(C)** The expression of the three *SNAT2* isoforms in normoxia (red bars) and hypoxia (0.1% O_2_, 48 hours; different blue bars for each isoform), measured by RPKM (Reads per kilo base per million mapped reads) by RNA-sequencing is shown in 4 breast cancer cell lines. **(E)** Schematic based on TMHMM prediction on the three SNAT2 isoforms identified by RNA-seq. The binding site for the SNAT2 antibody used for all the experiments recognizing the N-terminus of the protein is shown. (E) The relative expression of *SNAT2* isoform 1 and 2 by RT-qPCR is shown in hypoxia (0.1% O_2_, 48 hours; different blue bars for isoforms 1 and 2) compared to normoxia (red bars) in 4 breast cancer cell lines. *SNAT2* mRNA was normalized to the mean of *β*-actin and *RPL11*. Error bars, SD. Student t-test ***, p < 0.001; **, p < 0.01; *, p < 0.05; n = 3 for all the experiments

**Supplementary figure 2. SNAT2 is increased by hypoxia in a wide range of cancer cell lines. (A)**The relative expression of total *SNAT2* mRNA in SKBR3, T47D, MDA-MB-468 and HCC1806 breast cancer cell lines in normoxia and hypoxia. Results are obtained by using the mean of the Ct values of total *SNAT2* transcript after normalization to housekeeping genes (*β*-actin and *RPL11*). **(B)** The relative expression of total *SNAT2* in normoxia (red bars) and hypoxia (0.1% O2, 48 hours; blue bars) in breast cancer cell lines, prostate cancer cell lines, ovarian cancer cell lines and a colorectal cancer cell line is shown. *SNAT2* mRNA expression was normalized to the mean of *β-actin* and *RPL11*. **(C)** MCF7 cells were grown in normoxia or hypoxia (0.1% O_2_) for 24, 48 and 72 hours and total RNA was extracted and subjected to RT-qPCR analysis. *SNAT2* mRNA expression was normalized to the mean of *β*-actin and *RPL11*. **(D)** Immunoblot analysis of MCF7 lysates extracted from cells grown in parallel to those in (C). β-actin was used as a loading control. (E) The relative expression of total *SNAT2* and *CA9* mRNA in MCF7 and MCF7-HIF-1α-o in in normoxia and hypoxia is shown. Error bars, Student t-test. SD. ***, p < 0.001; ** p < 0.01; * p < 0.05; n = 3 for all the experiments.

**Supplementary figure 3. Bevacizumab decreases blood vessels and causes hypoxia and necrosis in breast cancer cell line xenografts (A-B)** *SNAT2* and *CA9* human mRNAs were increased in MCF7 (n=5) and MDA-MB-231 (n=5) xenografts treated with bevacizumab. mRNA expression was normalized to the mean of human *β*-actin and *RPL11*. **(C-D)** Representative H/E, CD31 (blood vessels), CA9 (hypoxia) immunostaining in MDA-MB-231 and MCF7 xenografts. Quantification with ImageJ of the staining for the selected antibodies in control (PBS) and treated (Bevacizumab) xenografts is shown on the lateral side. Scale bars: 200 μm; zoom, ×10. Non-parametric Mann-Whitney test, n=5 per group. Error bars, SD. ** p < 0.01, * p < 0.05.

**Supplementary figure 4. ERα and HIF-1α occupy the same genomic regions, but there are no addictive effects of estrogen supplementation under hypoxia in driving SNAT2 expression (A)** RNA-seq and ChIP-seq genome-browser view of occupancy (red box) of HIF-1α (top), HIF-2α (middle) and ERα (bottom) at the genomic coordinates of *SLC38A2* on chromosome 12 at high resolution. Peaks (red box) represent the areas where transcription factors interact with DNA. **(B)** RNA-seq and ChIP-seq genome-browser view of occupancy (red box) of HIF-1α (top), HIF-2α (middle) and ERα (bottom) at the genomic coordinates of *SLC38A1* and *SLC38A2* on chromosome 12. Peaks (red box) represent the areas where transcription factors interact with DNA. **(C)** MCF-7 cells were treated with 10nM E2 in 10% charcoal-stripped serum in normoxia and hypoxia (0.1% O_2_) for 30’, 2h, 4h, 12h, and 24 hours. The relative expression of total *SNAT2* in normoxia (red bars) and hypoxia is shown. *SNAT2* mRNA expression was normalized to the mean of *β*-actin and *RPL11*. Error bars, SD; n = 3 One-way ANOVA. ***, p < 0.001; ** p < 0.01; * p < 0.05; n = 3. **(D)** Immunoblotting of extracts from the same experiment is shown. β-actin is shown as a loading control.

**Supplementary figure 5. ERα and HIF-1α drive *SNAT2* expression independently (A)** RNA-seq and ChIP-seq genome-browser view of occupancy (red box) of HIF-1α (top), HIF-2α (middle) and ERα (bottom) at the genomic coordinates of *QSOX1* on chromosome 1. Peaks (red box) represent the areas where transcription factors interact with DNA **(B)** MCF7 cells were grown in charcoal stripped, phenol-free medium for 3 days and then incubated with or without fulvestrant (ICI 182,780; 10μM) and with or without 10 nM E_2_ in normoxia and hypoxia (0.1% O_2_) for 48 hours. *GAPDH, ALDOA* and *NEAT1* mRNAs were analyzed by RT-qPCR and normalized to 21% O_2_. mRNAs were normalized to the mean of *β*-actin and *RPL11*. Error bars, SD. One-way ANOVA. *** p < 0.001; **, p < 0.01; *, p < 0.05 n = 3. **(C)** Pathway map and GO processes enriched by MetaCore software of the 31 genes showing overlapping bindings sites for HIF-1α and ERα.

**Supplementary figure 6. SNAT2 overexpression promotes resistance to anti-estrogen treatment and glutamine deprivation (A)** Representative Western blots of SNAT2 knockdown in MCF7 cell line in normoxia and hypoxia. *SNAT2* knockdown reduced the level of phospho-S6 (p-S240/244-S6) in normoxia and phospo-mTOR and phospho-p70S6K in hypoxia. β-actin is shown as a loading control **(B)** SNAT2 and mTOR are partially colocalized on the Trans-Golgi network in MCF7 cells. Endogenous SNAT2 (green) is expressed in compartments that include the trans-Golgi network (TGN46; red). Although most mTOR (blue) is located elsewhere in the cytoplasm (probably late endosomes and lysosomes) (blue) in MCF7 cells, some mTOR staining also overlaps with the trans-Golgi network (red; indicated by arrows in merged images). DAPI (4', 6-diamidino-2-phenylindole) marks the nucleus. Scale bars are 5 □μm. **(C)** MCF7 cells were incubated in tissue culture medium depleted of the indicated amino acids in normoxia or hypoxia for 24 hours. Immunoblot analysis of SNAT2 is shown. β-actin is shown as a loading control; n=3 **(D)** 10^5^ MCF7 cells either with or without *SNAT2* knockdown were seeded in normal, glutamine-free or glucose-free medium for 3 days in normoxia (red) and hypoxia (1% O_2_; blue), and final cell number counted; n=3. One-way ANOVA (E) Representative Western blots and confocal imaging of SNAT2 (green) overexpression in MCF7 (MCF7-SNAT2-o) cell line versus parental control cells. β-actin is shown as a loading control. DAPI (blue) Scale bars are 10 μm **(F)** 10^5^ cells were seeded after 3d in charcoal-stripped medium and were treated with or without E_2_ (10nM), fulvestrant (ICI 182,780; 10uM) in MCF7 parental and MCF7-SNAT2-o cells for 5 days in charcoal-stripped medium in normoxia and hypoxia (1% O_2_); n=4. One-way ANOVA **(G)** Representative graph of the effect of SNAT2 overexpression (red) versus control cells (black) in MCF7 spheroid growth in different growth medium (normal DMEM (5mM), low glucose (1mM), low glutamine (1mM)). Two-way ANOVA. Error bars, SD. *\*\*\**, P < 0.001; **, P < 0.01; *, P < 0.05 for all the experiments.

**Supplementary figure 7. SNAT2 levels correlate with HIF-1α but not c-Myc expression in breast cancer patients *in vivo*. (A)** Correlation heatmap of *SNAT2* and other genes part of the hypoxia signature, AA transporters, *HIF-1α*, and *ATF4, SNAT2* co-expresses (black box) with HIF-1α and other genes (*ATF4, HK2, MCTS1, P4HA1*, CORO1C)(black box) previously identified as part of the hypoxia signature in breast cancer patients. Each square represents the Pearson correlation (r) between a pair of genes, calculated using microarray expression data from the Metabric cohort. Red colors indicate a high gene–gene correlation while the opposite is seen for the blue. **(B)** *Myc* copy number and *SNAT2* mRNA expression do not correlate in the Metabric breast cancer cohort.

